# Voluntary action sharpens sensory prediction and facilitates neural processing of contingent sensory stimuli

**DOI:** 10.1101/2024.12.03.626576

**Authors:** Edward Ody, Tilo Kircher, Benjamin Straube, Yifei He

**Affiliations:** Department of Psychiatry and Psychotherapy, University of Marburg, Rudolf Bultmann-Strasse 8, 35039 Marburg, Germany

## Abstract

Self- and externally generated sensations differ in sensory responses and motor preparation. However, mechanistic evidence linking the two is scarce. Here, participants made active (self-initiated) or passive (finger moved by electromagnet) movements that triggered a bimodal auditory/visual stimulus. These were followed, in a block-wise manner, by a unimodal auditory/visual comparison stimulus, and participants judged which stimulus was brighter or louder. Motor preparation ERPs encoded task modality and movement, while sensory ERPs showed reduced task differences for active, demonstrating sensory suppression. Next, we decoded task modality within active and passive. During motor preparation, the active condition showed higher accuracy, reflecting enhanced predictive processes. In the sensory perception period, accuracy was higher in the passive condition, mirroring previous reports of reduced sensory responses to self-generated stimuli. Temporal generalisation showed pattern similarities in the alpha band amplitude between the motor preparation and the stimulus perception windows. This suggests that alpha oscillations may encode sensory predictions generated during motor preparation. Our findings provide mechanistic evidence of action-effect prediction during voluntary actions.

## Introduction

Distinguishing between self- and externally generated sensations is fundamental to having meaningful interactions with the external world, maintaining a sense of agency, and enhancing the precision of movements ^1^. This distinction is facilitated by an internal forward model ^2,3^. When preparing to execute an action, the brain prepares a prediction of the expected sensory consequences of that action. Incoming sensory input is then compared to what was predicted and by observing the discrepancy between the two, the brain can determine whether that input was self- or externally generated. Though these two processes (prediction and comparison) are assumed to arise from a cohesive system, they have almost exclusively been studied in isolation, with evidence linking them together largely remaining correlational ^4–6^.

Previous EEG studies have demonstrated both sides of this action-perception loop. In the motor preparation time window, ERPs ^7–10^ and multivariate pattern analysis (MVPA) of EEG ^9,11^ encode the action’s sensory consequences, demonstrating action-effect prediction. Self- and externally generated sensations are subsequently processed differently, as shown in early sensory (N1 and P2) ERP components ^12–15^. Earlier studies typically showed a reduction in sensory responses for self-generated action consequences ^12,15–18^ (sensory suppression). However, more recent studies have also demonstrated perceptual enhancement ^19–22^. This has led to competing theories regarding how sensory modulation arises. The suppression results were typically attributed to a cancellation account ^16^, in which self-generated and predictable sensations are neurally and perceptually attenuated ^23–25^. An alternative Bayesian framework (the sharpening account) reconciles the enhancement findings ^26–29^. It posits that neural activity in sensory areas is tuned toward what is expected based on the accumulation of previous evidence. Accordingly, sensations that are predicted by voluntary action should be up-weighted.

Both the cancellation and the sharpening accounts predict different effects for sensory perception but converge on the idea that action preparation encodes sensory prediction. However, empirical evidence showing how the action’s motor preparation relates to subsequent sensory perception is missing. Here, with a novel design, we examined motor preparation, sensory perception, and the neural mechanism that may mediate between them. We embedded an intermodal attention manipulation ^30–33^ into a contingency paradigm (Figure 1A) ^24^. Participants made active (voluntary) or passive (finger moved by device) movements that led to a bimodal (auditory and visual) stimulus (tone via headphones and visual solid disc on a screen), always presented at the same intensity. Using passive finger movements in the contingency paradigm provides optimal control over differences in temporal prediction and tactile feedback ^14^ between the active and passive conditions. The movement-contingent stimulus was followed, in a block-wise manner, by a unimodal visual or auditory comparison stimulus. Participants judged whether the movement-contingent stimulus or comparison stimulus in each trial was louder/brighter (depending on the task condition), directing their attention to one or other modality. We expected the modality effect to differ between the movement conditions, driven by additional predictive processes in the active condition. Using a combination of traditional univariate ERPs and multivariate pattern analysis (MVPA), we compared task modality (auditory vs visual) differences between the active and the passive movement conditions. We then tested for mechanistic evidence linking the motor preparation and sensory processing periods together.

**Figure 1.**
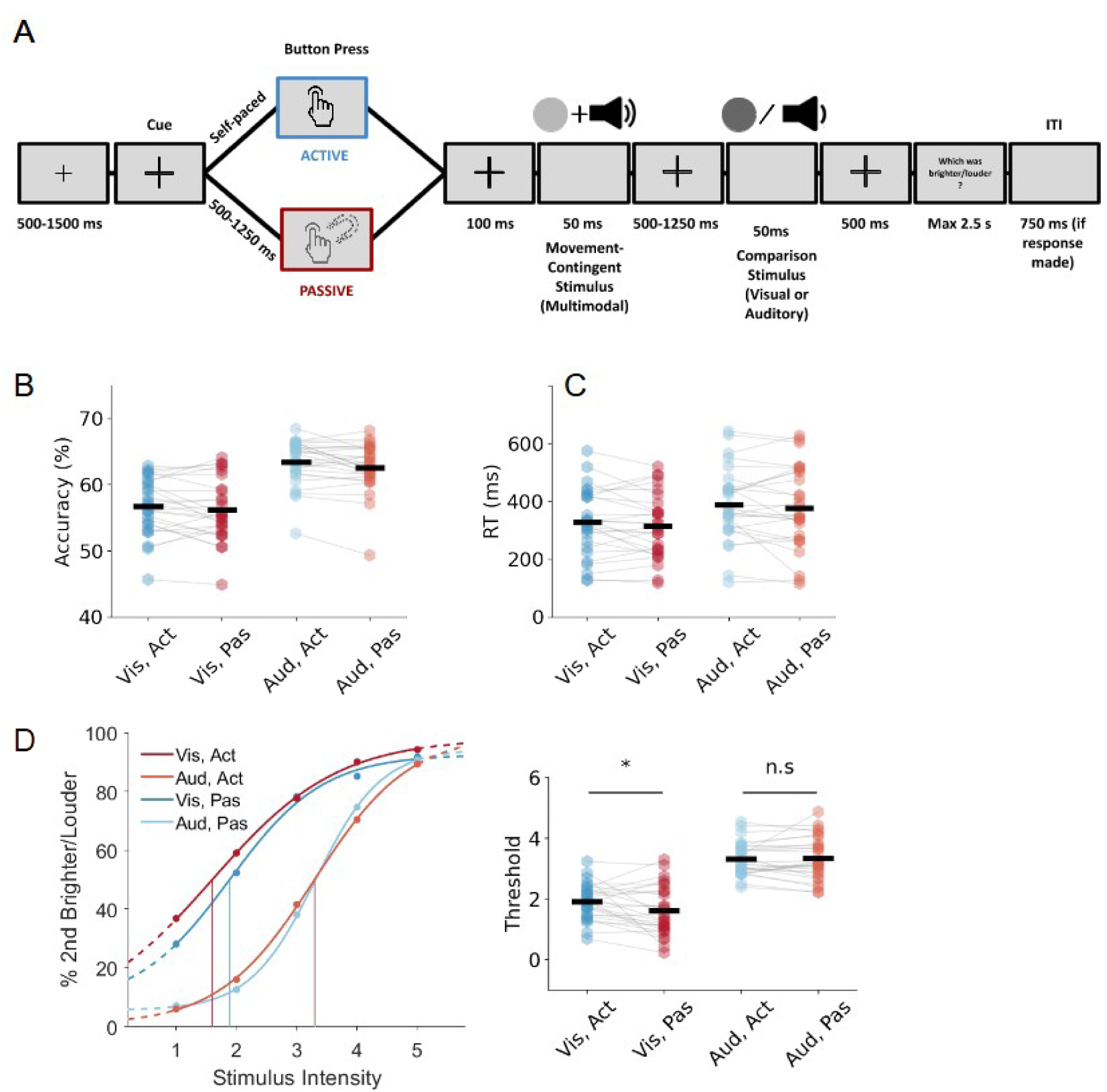
Paradigm and behavioural results. (A): The trial structure. Participants made active (self-generated) or passive (finger moved by electromagnet device) button presses which were followed by a bimodal (auditory and visual) movement-contingent stimulus. This was then followed by a unimodal auditory *or* visual comparison stimulus and participants had to judge whether the movement-contingent or comparison stimulus in that trial was brighter/louder. (B, C) Accuracy and response times for responding to the intensity judgement task. (D) Intensity judgement task results as revealed by psychometric functions Left panel shows average thresholds (for illustrative purposes only. Statistics were performed on thresholds derived individually for each participant).

## Results

### Behavioural Results

Participants performed an intensity judgement task (Figure 1A) where they judged whether the first (movement contingent) or second (comparison) stimulus on each trial was brighter or louder (depending on whether the block was visual or auditory). Psychometric functions were fitted to the data (Figure 1D) to derive points of subjective equality (thresholds) for each participant and each condition. In the visual task, passive movements had significantly lower thresholds than active ones (*t*(1,26) = 2.3, *p* = .030), suggesting that the movement-contingent stimuli (those presented directly after the button press) were perceived as *less* intense in the passive condition compared to the active condition.

According to the cancellation theory, it would be expected that self-initiated sensations are perceived as less intense ^34,35^. This result goes against that hypothesis but supports the notion that self- and externally produced sensations are perceived differently. Participants responded faster in the visual condition (*F*(1,26 = 5.98, *p* = .022, 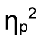 = 0.19) but were less accurate (*F*(1,26 = 53.79, *p* < .001, 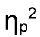 = 0.67) (Figure 1B, 1C). For response times and accuracy, none of the other effects was significant (all *p* > .05). Detailed statistics are presented in the supplement (S1).

### RP encodes movement and task modality, while LRP only encodes movement

We first examined traditional ERPs in the motor preparation (RP and LRP) and stimulus perception (N1-P2 peak-to-peak) time windows, reflecting the two sides of the action-perception loop. Beginning with the pre-motor time window, we investigated typical motor preparation ERPs, Readiness Potential (RP), and Lateralized Readiness Potential (LRP). RP is a slow negative wave related to higher-level movement preparation. It has been shown to encode the type of movement ^9^, its sensory consequences ^7–10^, and task-related information such as action-effect contingency ^9^. Therefore, we first investigated whether the RP encoded task modality using cluster-based permutation statistics (see Methods). The test revealed a significant cluster beginning around 200 ms before the button press. Within this window, we expected larger differences between visual and auditory task modality in the active condition than in the passive condition, due to additional motor-related predictive processes associated with self-initiated movements ^7–9^. The results showed that RP encoded the movement type *F*(1, 27) = 16.54, *p* < .001, η_p_ ^2^ = 0.38 and the task modality *F*(1, 27) = 7.63, *p* = .01, η_p_ ^2^ = 0.22. However, the interaction between movement and task modality was not significant (*F*(1, 27) = 0.62, *p* = .437, η_p_ ^2^ = 0.02), suggesting that the modality effect was not modulated by the movement type (Figure 2A-C).

**Figure 2.**
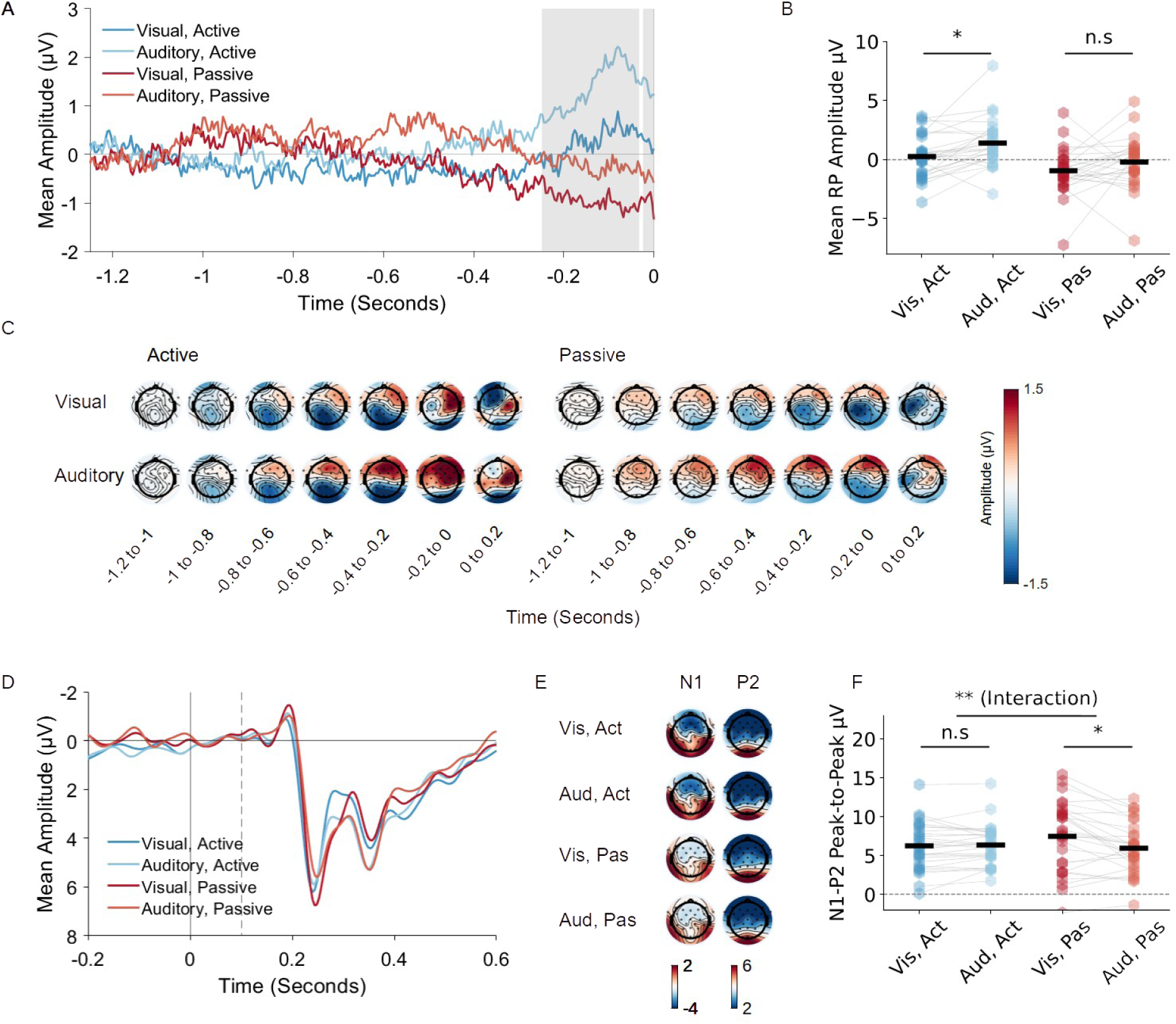
Event-related potential results. (A): Readiness potential. Average of electrodes Cz, C3, and C4, time- locked to the button press (t = 0). The grey shaded area shows time periods where the permutation test (auditory vs. visual, across movements) identified significant clusters. (B) Mean amplitudes averaged across all significant time points identified in the permutation test. (C) Topographical plots for all experimental conditions in 200 ms bins for the 1200 ms time period preceding the button press. (D) Visual ERPs averaged across electrodes Oz, O1, and O2, time-locked to the button press (0), with stimulus presentation at 0.1 seconds (dashed line). Baseline period 0 to 0.1 seconds (interval between button press and stimulus). (E) Topographical plots for the N1 (80-104 ms) and P2 (132-156 ms) peaks. (F) Mean N1-P2 peak-to-peak amplitude for the four conditions.

Secondly, we investigated the LRP, derived by subtracting activity at one side of the scalp (C3) from activity at the other side (C4). Due to its lateralized nature, LRP is considered to reflect lower-level motor-specific preparation for action execution, originating in the primary motor cortex (M1). Most studies suggest that it does not encode action-effect contingency ^9,36^. Consistent with these reports, LRP was sensitive only to movement type (*F*(1, 27) = 24.48, *p* < .001, η_p_ ^2^ = 0.48), showing more negative amplitudes in the active condition (estimated marginal mean; EMM = -2.06 μV) than the passive condition (EMM = - 0.97 μV). The main effect of task modality (*F*(1, 27) = 1.97, *p* = .172, η_p_ ^2^ = 0.07) and the interaction between movement and task modality (*F*(1, 27) = 3.02, *p* = .094, η_p_ ^2^ = 0.1) were not significant (Figure S2AB).

### In the sensory ERPs, the modality effect was reduced in the active movement condition

Regarding the primary sensory ERPs, we focused on visual ERP response difference (O1, O2, Oz) between task modality (intermodal effect ^30–33^), and expected a reduced effect in the active condition following prior literature ^12,13,15,18^. N1-P2 peak-to-peak amplitude was analysed to provide a better signal-to-noise ratio. There was a significant interaction between movement and task modality (*F*(1, 27) = 11.84, *p* = .002, 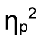 = 0.31). N1-P2 peak-peak amplitudes were significantly larger in the visual task than the auditory task for passive movements (*t* = 4, *p* = .001, Bonferroni-corrected) but there was no significant difference between visual and auditory for active movements (*t* = -0.32 *p* > .999, Bonferroni-corrected). In other words, the intermodal attention effect was reduced for the active movements, showing an active suppression (Figure 2D-F). The main effect of task modality was significant (*F*(1,27) = 5.6, *p* = .025, η_p_ ^2^ = 0.17), with higher amplitudes for visual (EMM = 6.86) than auditory (EMM = 6.17). The main effect of movement was not significant (*F*(1,27) = 0.92, *p* = .35, η_p_ ^2^ = 0.03). Due to additional noise from the internal mechanism in the passive condition, the auditory ERP results were likely confounded (Supplement S1). Auditory results and details are given in the Supplement S2.

### Active movement showed enhanced modality decoding in motor preparation but decreased decoding in stimulus perception

We performed a series of MVPA analyses to investigate how motor prediction and sensory perception are encoded in patterns of EEG activity across the scalp and to examine how the two time windows are related. First, we trained a classifier (linear discriminant analysis; LDA) to discriminate task modality (visual or auditory) at each time point from 1000 ms before the button press to 600 ms afterwards (baseline corrected at -1250 to -1000 ms), separately for active and passive movements. We first applied this analysis to the broadband signal. In the motor preparation window (-400 to 0 ms), both movement conditions showed an above-chance, ramping decoding (pmax < .004, tmin = 3.15, Figure 3A), with significantly higher accuracy in the active than the passive condition (p < .002, t = 3.37, Figure 3B, left).

**Figure 3.**
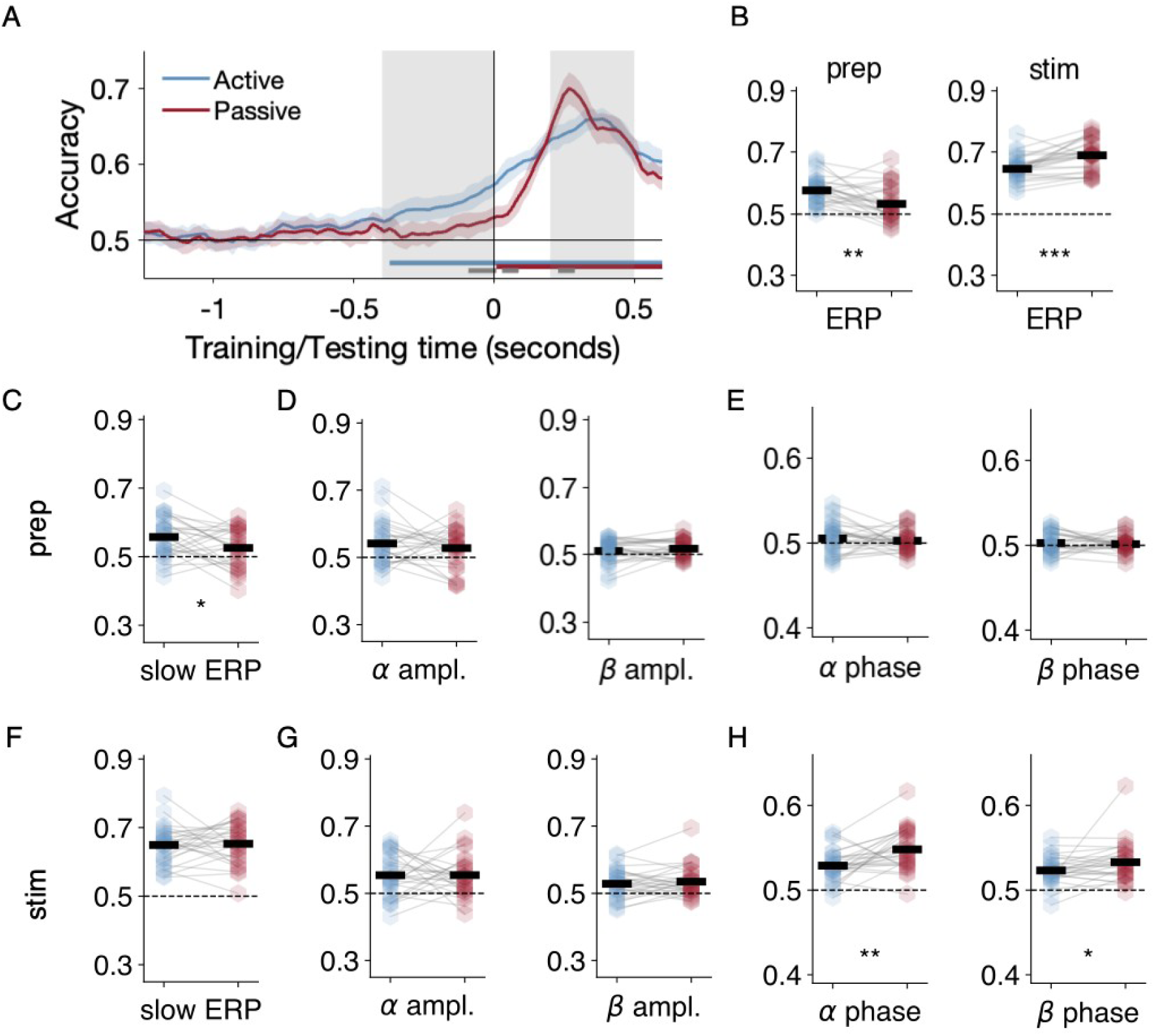
Decoding task modality (auditory vs. visual) across time for the active (blue) and the passive (red) conditions. (A): Decoding accuracy time series (means and standard errors) based on the broadband ERPs. Above-chance decoding was already observed within both the motor preparation (prep., -0.4 to 0 seconds) and the stimulus perception (stim., 0.2 to 0.5 seconds, corresponds to 0.1 to 0.4 seconds after stimulus onset) time window (time of interests, TOIs), as highlighted by the light grey patches. Horizontal thick lines of respective colours indicate the time windows where the accuracy of both conditions was significantly above chance (cluster p < .05, corrected). The grey horizontal thick line illustrates the time windows where the accuracy of both conditions significantly differ (cluster p < .05, corrected). Time zero marks the onset of the button press. Audiovisual stimulus onsets at 0.1 seconds. (B-H): Decoding accuracy for individual participants (blue and red hexagons) averaged within the motor preparation and stimulus perception TOIs. B: Decoding accuracy based on broadband ERPs within the motor preparation and stimulus perception windows suggests significantly higher decoding for the active movement during motor preparation, whereas during stimulus perception, task modality was better decoded in the passive condition. (C-E): within the motor preparation window, decoding accuracy based on the slow ERPs (C), the alpha and beta amplitudes (D), and the alpha and beta instantaneous phase (E). (F-H): within the stimulus perception window, decoding accuracy based on the slow ERPs (F), the alpha and beta amplitudes (G), and the alpha and beta instantaneous phase (H). For B-H, Horizontal black bars indicate the mean across all participants. The means significantly above chance are marked as solid; when they do not differ significantly from chance they are marked as dotted. Asterisks indicate statistical significance of dependent- sample t-tests between the active and passive conditions, p < .001: ***, p < .01: **. p < .05: *.

This suggests that there may be enhanced predictive processes related to the action outcome present in the active condition, resulting in larger EEG pattern differences between modalities available to the classifier and ultimately, higher decoding accuracy. The ramp-up pattern also suggests that this prediction enhances over time until the action (button press) is executed. In the stimulus perception window (200 to 500 ms after button press, corresponding to 100 to 400 ms post-stimulus onset), task modality was also decodable in both movement conditions (pmax < .001, tmin = 19.29, Figure 3A). However, in contrast to the motor preparation window, here, the passive condition showed higher decoding accuracy than the active condition (p < .001, t = 4.74, Figure 3B, right). Notably, this effect remains stable if we apply a 100 ms pre-stimulus baseline (0 to 100 ms relative to button press), as reported in Figure S3. We also compared the decoding time series between movements based on EEG signals at the second (comparison) stimulus. Importantly, despite clear decodability between the two unimodal inputs, we observed almost completely overlapping decoding time series between both movement conditions (Figure S4). The reduced modality decoding suggests dampened sensory responses in the active condition, potentially reflecting the same modulation of sensory processing typically seen in ERPs. More importantly, the specificity of this decoding difference at the first, but not the second stimulus, suggests that this effect is movement-contingent. We followed up the broad-band decoding with band-specific decoding analyses, focusing on the slow ERP ^37^ (< 6Hz), and the amplitudes and instantaneous phase within the alpha (8–12Hz) and the beta (13–30Hz) bands ^38–40^, Figure 3C-H. The decoding time series of these signals is reported in Figure S5. In the motor preparation window, only the slow ERPs showed significant decoding of task modality in both active and passive conditions (pmax < .025, tmin = 2.26) as well as significantly higher accuracy for the active condition (p < .036, t = 2.21, Figure 3C). This suggests that the pre-stimulus active/passive difference in predicting task modality was primarily driven by the slow ERPs. In the stimulus perception window, however, we observed significantly higher decoding accuracy in the passive condition, only in analyses based on instantaneous phase within the alpha (p < .002, t = 3.49) and the beta bands (p < .047, t = 2.09), as in Figure 3H. These results suggest that the task-perceptual difference between movements was predominantly phase-locked and therefore driven by the sensory ERPs. We also carried out analogous analyses in the delta and theta frequency bands, and reported corresponding decoding time series and their extracted individual (and mean) accuracies in Figures S5 and S6.

### The role of the alpha oscillations in forward model mechanisms

Finally, we tested for evidence of potential neural links between motor preparation and sensory perception. We used temporal generalisation (training the classifier on each time point and testing on every other time point) to test if neural signals share any pattern similarities across the two time windows ^41^. When decoding using the broadband ERP, we did not observe any significant temporal generalisation between the two time windows in either movement type (Figure 4AB). However, in the analysis based on alpha amplitude, we observed significant above-chance temporal generalisation in the active condition (p = .013, t = 2.66, Figure 4C, left) and not in the passive condition (p = .438, t = 0.78, Figure 4C, right). However, the difference in accuracy between the movements was not significant (p = .176, t = 1.39, Figure 4D). These findings suggest that, in the active condition, alpha amplitude’s neural pattern is shared between the motor preparation and stimulus perception windows. We also performed this analysis with other signals but observed no significant temporal generalisation (Figure S7). Based on this finding, we further tested if task decoding performance in the motor preparation window relates to that of the stimulus perception window on a by-subject level (Figure 4E). We observed a significant positive correlation within the active condition (Pearson’s *r* = 0.47, *p* = .012) but not the passive condition (Pearson’s *r* = 0.31, *p* = .111). Concatenating both movement conditions, this correlation was also positive and significant (Pearson’s, *r* = 0.39, *p* = .003). Lastly, we tested if active and passive conditions share neural patterns in either motor preparation or stimulus perception time windows with cross-decoding analyses, by training the classifiers from one pair of conditions and testing them on the other pair (Figure S8). Results from cross-decoding showed that, during motor preparation, alpha amplitude patterns between both movements were cross-decodable. However, for stimulus perception, active and passive movements were decodable based on predominantly phase-locked signals. To summarise, in a contingency paradigm, the prediction for the modality of an action’s sensory consequences may be encoded within the alpha band.

**Figure 4.**
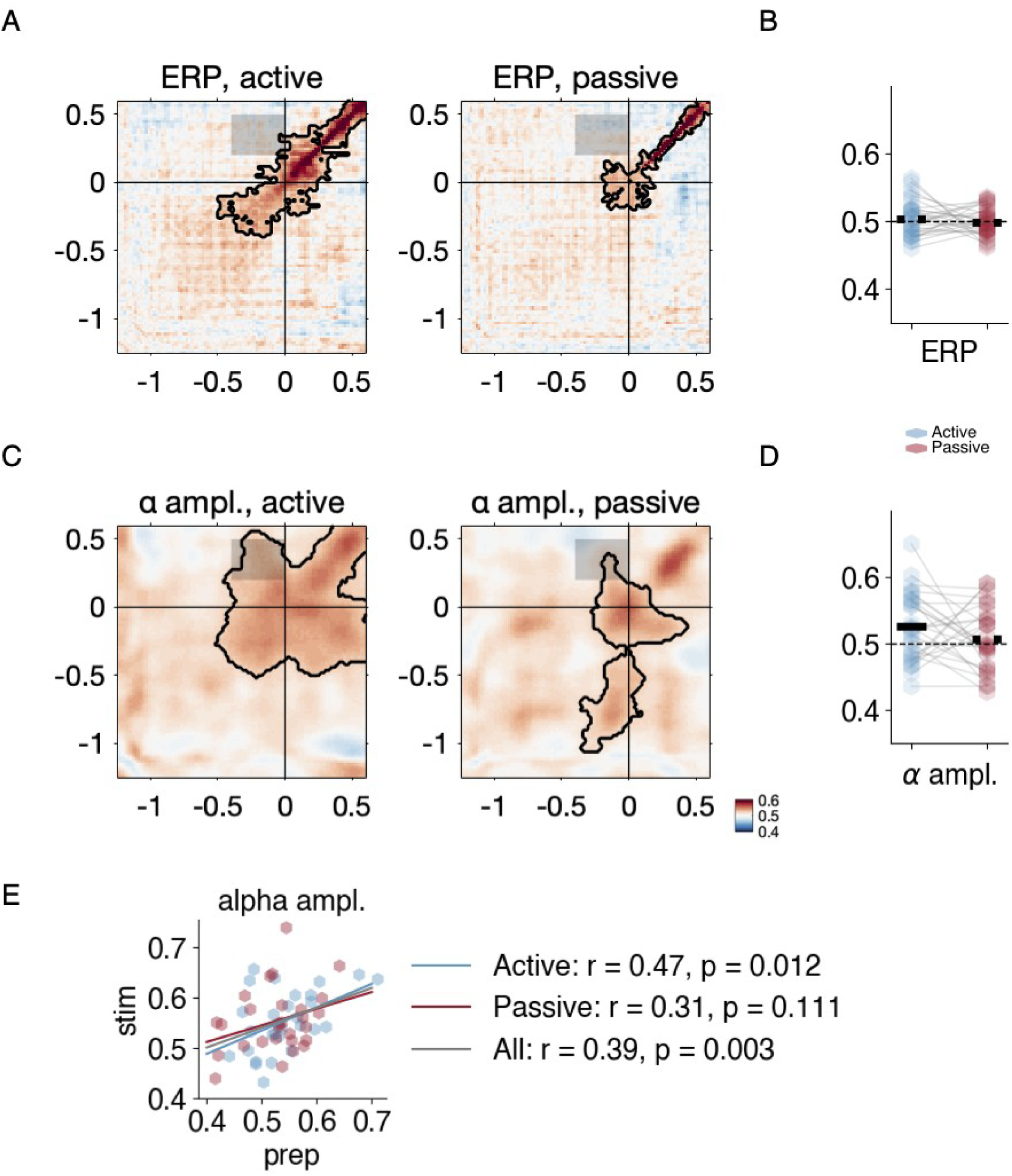
(A): Temporal generalisation based on the broad-band ERPs for the active (left) and the passive (right) conditions. We observed no significant temporal generalisation (highlighted by the black contour) training from the motor preparation window (-0.4 to 0 seconds) and testing on the stimulus perception window (0.2 to 0.5 seconds, corresponds to 0.1 to 0.4 seconds after stimulus onset), as shown within the grey patch. (B): Extracted averaged temporal generalisation accuracy showed no significant effects in both conditions. (C): Temporal generalisation based on alpha amplitude showed significant decoding training from motor preparation and testing on stimulus perception. (D): Extracted averaged temporal generalisation accuracy for both active and passive conditions suggests significant decoding only for the active condition (blue) but not the passive condition (red), without significant difference between both conditions. For A&C, the black contour indicates training-testing time point combinations with above-chance decoding, corrected with a cluster-based permutation test. For B&D, Horizontal black bars indicate the mean across all participants. The means significantly above chance are marked as solid; when they do not differ significantly from chance they are marked as dotted. Asterisks indicate statistical significance of dependent-sample t-tests between the active and passive conditions, p < .001: ***, p < .01: **. p < .05: *. (E): by-correlation between the averaged decoding performance within the motor preparation window (x-axis) and the stimulus perception window (y-axis) for both the active (blue) and the passive (red) conditions.

## Discussion

The current study aimed to examine the mechanisms underlying motor prediction. We used both univariate ERP methods and MVPA to investigate the neural activity associated with action-effect prediction during the motor preparation phase and with the sensory processing of the action’s sensory consequences. Participants made active (self-initiated) or passive (finger moved by electromagnet) movements that triggered a movement contingent bimodal (auditory/visual) stimulus. This was followed, block-wise, by an unimodal comparison auditory *or* visual stimulus, with varying intensity levels. Participants had to judge which stimulus in each trial was brighter/louder, directing attention to one modality or the other.

First, we found that conventional motor-preparatory ERPs did not encode sensory prediction specific to voluntary movement. Readiness potentials were sensitive to movement type and task modality but there was no evidence of an interaction between the two factors. In other words, RP encoded the upcoming task modality but it appeared to do so equally well in both the active and passive movement conditions. This goes against our initial hypothesis that the modality effect would differ between the movements, due to additional predictive processes in the active condition. Previous studies show that RP contains a motor-specific prediction that cannot be explained by mere anticipation of an upcoming stimulus ^7–9^.

Conversely, the current result suggests that RP captured more general predictive processes that were not purely motor-specific. However, this univariate ERP analysis may have simply been too restricted to capture the interaction effect, as the more sensitive whole-scalp MVPA (see below) was able to identify motor-related differences in the modality effects. Unlike RP, LRP was sensitive only to the movement type, showing a sharp deflection of approximately 200 ms before the button press in the active conditions. This result is consistent with the idea that LRP reflects lower-level motor-specific preparation for action execution ^9,36,42^.

Together, the pre-motor ERP results support the notion of a functional dissociation between RP and LRP, with the former being associated with higher-level movement preparation (such as the decision to move, movement velocity, movement trajectory, and prediction for action consequences, as in the current design).

However, when we looked beyond these univariate motor-preparatory ERPs, with MVPA, we found that EEG signals prior to the button press did encode action-specific sensory prediction. In the motor preparation time window, the classifier showed a ramping decoding pattern in both active and passive movements, with above-chance decoding in the 400 ms before the button press. This finding is consistent with the RP results, showing that the prediction for the task modality is already encoded in the pre-motor period and that this prediction strengthens over time until motor execution. However, while RPs did not show an interaction between movement and task modality, MVPA revealed differences between the movement conditions. Active movements showed significantly higher decoding accuracy than passive movements, suggesting a sharpened representation of the action’s sensory consequences in the active condition, as more pattern information relating to the task condition was available to the classifier. Furthermore, while the interaction was not detected in the univariate analysis, frequency-band-specific decoding showed a significant movement difference in the slow ERP (below 6 hz), but not in the delta/theta bands. This might suggest that slow waves were implicated in encoding the movement differences ^43,44^. We have previously shown that MVPA can reveal more subtle differences between conditions by taking into account the pattern of activity across the whole scalp ^9^. These results show that action-effect prediction may be spread more widely across different electrodes and that the effects are not necessarily captured entirely by univariate ERPs. Taken together, our pre- motor results not only suggest how action-specific sensory prediction unfolds over time but also show that enhanced prediction from the active movement is carried out in slow motor-preparatory EEGs. Overall, this result supports the idea that predicted sensory consequences are generated gradually during movement preparation ^8,9^.

Regarding sensory perception, the ERP and MVPA results both align better with cancellation theories. When looking at the ERPs associated with sensory perception, we observed an interaction between movement type and task modality. In the passive condition, there was a significant difference between the visual and auditory modalities, with smaller N1-P2 peak-to-peak amplitudes in the auditory condition. There was no evidence of a difference between the task modalities in the active condition. This result could be interpreted under the cancellation account of motor control. According to these theories, sensory responses for predictable, self-generated sensations are reduced or cancelled out, allowing attentional resources to be allocated to important, external events. The current result suggests that the sensory responses were reduced in the active condition, leading to more similar processing between the auditory and visual conditions. While intermodal attention effects have been studied before ^30–33^, to our knowledge, this is the first evidence that the effect is modulated by action. This finding corroborates prior studies that demonstrated a reduction in early sensory ERP component amplitudes for self-generated sensations ^12,14,15,17,18,36,45^ (but see ^19,20,29,46,47^). Nevertheless, the current results support the cancellation hypothesis and a reduction in sensory processing of self-generated action consequences.

Importantly, when approaching sensory perception with MVPA, although the emerging sharpening theories would predict enhanced modality decoding for the active movement during action-outcome processing, our results revealed the opposite pattern. In the sensory perception window, classification accuracy was high, reaching approximately 65-70%, showing that task modality could be decoded given identical bimodal stimulation. Importantly, accuracy was higher in the passive condition than in the active condition. This decoding difference was only observable from EEG time-locked to the movement-contingent but not the comparison stimulus, suggesting that the effect is specific to action contingency. This finding might reflect a reduction in the sensory responses in the active condition, as observed in the ERPs, as there would be reduced EEG pattern difference between modalities, resulting in lower accuracy. This argument is further supported by the narrow- band analysis, which showed that the instantaneous phase in the alpha and beta bands was the most prominent signal to reveal the active vs. passive differences. Notably, these results are inconsistent with previous MRI studies ^26,29^. For example, Yon et al ^29^ compared index and little finger movements that lead to congruent or incongruent finger movements of an avatar hand. Classification accuracy for decoding the stimulus finger movement was higher for congruent than incongruent actions. Kok et al ^26^ showed that gratings with an orientation that could be predicted based on an auditory cue had sharper representations in the primary visual cortex. This suggests that the expectation of sensory consequences sharpened the representation of the action’s effect, contradicting the current result. However, there are marked differences between the methods of those studies and the current one. The previous studies manipulated expectancy differently from the current study, using congruency or proportional methods. Conversely, in the current study, we manipulated motor prediction using active and passive movements. Secondly, the factors being decoded in the previous studies were more fine-grained than the current study, which tested the broader difference between sensory modalities. Future studies could shed further light on the issue of attenuation vs enhancement by investigating how these factors interact, for example varying the proportion of predictable stimuli triggered by active and passive movements. Taken together, the evidence from both time windows suggests that active movements sharpen sensory prediction during motor preparation and subsequently facilitate sensory processing by reducing neural differences between modalities.

The second core goal of the study was to investigate the neural link between motor preparation and action feedback processing. It has been suggested that perceptual predictions may be encoded within neural oscillations, particularly in the alpha and beta bands ^38,48–51^. Notably, however, prior studies have only shown correlational evidence linking motor prediction to sensory perception of self- and externally generated sensations ^4,13^.

Here, our results point to a facilitatory role of alpha-band oscillations in sensorimotor integration. Alpha amplitude (but not broadband ERP) showed patterns of neural activity that were generalised between the motor preparation and sensory perception windows. A recent study by Hetenyi et al ^38^ showed that an upcoming shape stimulus that was predicted by an auditory cue was represented in pre-stimulus alpha oscillations. In a similar vein, our current result suggests that motor prediction may also be mediated by alpha oscillations. Qualitatively, the activity was greater in the active condition, although there was not a significant difference between the movement types. However, cross-decoding showed that the neural pattern was similar between movements (Figure S7). There was also a significant positive correlation between decoding accuracy based on alpha amplitude in the motor preparation and sensory perception time windows. Taken together, these results provide preliminary evidence that alpha-band activity carries motor prediction from action preparation to sensory perception.

In the behavioural data, discrimination thresholds in the visual task were higher in the active condition than the passive condition, suggesting enhanced perception for the active condition. According to the cancellation theory, sensory responses for self-generated sensations are attenuated or cancelled out, as the motor system has prepared the sensory cortices for the expected action feedback. According to this theory, lower thresholds are expected in the active condition, as this would indicate that perception of the stimulus triggered by the button press was dampened. The current result contrasts previous studies showing reduced perceptual thresholds for self-generated sensations ^35,52^ and also suggests a different effect from the reduced sensory ERPs and lower decoding accuracy in the EEG results. However, some studies have shown enhanced detection for active conditions ^21,53^. It has been shown that the overall stimulus intensity (for example, near-threshold or suprathreshold) can affect whether self-generated sensations are suppressed or enhanced ^54,55^, so the current effect might be related to low luminance/volume of the stimuli.

Furthermore, neural responses associated with self-generated sensations can be attenuated even when behavioural responses are enhanced ^56^.

A long-standing line of literature shows that self-initiated movements are associated with a prediction for their sensory consequences that occurs before the movement, and that a modulation of the sensory responses is associated with the action outcome of the movement. Here, by employing a novel paradigm that manipulates intermodal attention in a contingency paradigm, with both univariate ERPs and MVPA, we went beyond this literature and provided cohesive evidence that suggests sharpened action-effect prediction before the movement execution and facilitated sensory processing after the stimulus presentation. We also observed a robust pattern generalization effect, demonstrating the potential role of alpha oscillations in carrying predictions between motor preparation and sensory perception. Together, these data provided fine-grained mechanistic evidence of action-effect prediction.

In this study, we investigated both processes using an intermodal attention paradigm embedded within a contingency paradigm. In both time windows, we demonstrated active movement preference, with sharpened action-effect prediction before the movement execution and facilitated sensory processing after the stimulus presentation. These results were evident from both the univariate ERPs and from the MVPA, with the latter providing more nuanced details. We have also demonstrated the role of alpha oscillations in carrying predictions between motor preparation and sensory perception. Overall, our findings provide fine-grained mechanistic evidence of action-effect prediction during voluntary actions.

## Method

### Participants

Twenty-eight participants (18 female) took part, with ages ranging between 19 and 31 (M = 23.8, SD = 3.4). Participants received a €30 inconvenience allowance for taking part. The local ethics committee approved the study protocol in accordance with the Declaration of Helsinki (except for pre-registration) ^57^, and all participants provided written informed consent. By self-reported, participants were right-handed, had normal or corrected-to-normal hearing and vision, no history of mental illness, no history of drug or alcohol abuse, no history of serious brain injury, and no first-degree relatives with schizophrenia.

Participants were naïve to the purpose of the experiment. Due to a technical error with the presentation software, one participant’s behavioural data were missing. This participant was included in EEG-only analyses but excluded from analyses including behavioural data.

### Task and Procedure

The experiment was run in a semi-darkened room on a 19” 60 Hz computer monitor. Participants sat with their right index finger resting on a button pad and attached to it with a piece of elastic. The left hand was placed on the computer keyboard. Pink noise was through headphones at 45 dB throughout the experiment to mask the sound of the button press. Participants first completed a training session to familiarise them with the task. This consisted of 5 blocks of 5 trials each. The EEG cap was then fitted prior to starting the main experimental and control blocks.

Participants completed an intensity judgement task where they were asked to indicate which of two successively presented stimuli was brighter or louder. The stimuli could be visual (grey discs) or auditory (sine wave tones), presented in separate blocks. However, critically, the movement-contingent stimulus in each trial was always presented multimodally, with visual and auditory stimuli together. Stimuli were triggered by active or passive button presses. Active button presses were self-initiated while passive button presses were produced using an external device that pulled the finger down.

The experiment consisted of 2 experimental blocks and 2 control blocks. Each experimental block contained 200 trials which were further divided into active and passive mini blocks containing 25 trials (making a total of 4 active and 4 passive mini blocks per experimental block). This meant that each stimulus level of the comparison stimulus was presented 20 times. Participants always completed both of the auditory blocks together and both of the visual blocks together. Counterbalancing was achieved by varying the order of presentation (visual or auditory first) and the starting action in each block (active or passive movement). The data were collected during a session that also included multimodal blocks which were not the focus of the present study. These blocks were interspersed with the described blocks. All visual blocks and all auditory blocks were always presented together. However, the order of unimodal and multimodal blocks, the order of auditory and visual blocks and the order of active and passive trials within the blocks was counterbalanced. 50% of participants started with visual and 50% started with auditory. 50% started with unimodal while 50% started with multimodal. 15 participants started with active trials while 13 started with passive.

Each trial began with a fixation cross. After a random duration between 500 and 1500 ms (in 250 ms steps), the cross enlarged, acting as a cue for the button press. In active trials, participants pressed the button at their own pace after this cue. In passive trials, the button moved, pulling the finger down, after a random interval between 500 and 1250 ms (in 83 ms steps). To control for temporal prediction, there was a fixed 100 ms interval after the button press before the first stimulus (movement-contingent stimulus) was presented. This stimulus consisted of both an auditory stimulus and a visual stimulus together, presented at a fixed intensity. After a random duration between 500 and 1250 ms (in 250 ms steps), a second (comparison) stimulus was presented. This stimulus was always presented unimodally at one of five different intensities, two lower, two higher and one identical to the stimulus of interest. All stimuli were presented for 50 ms. After a 500 ms pause, the question ’Welcher war heller?’ (’Which was brighter?’, in German) was shown on visual trials and ’Welcher war lauter?’ (’Which was louder?’) on auditory trials. Participants pressed ’v’ on the keyboard to indicate ’first stimulus’ (movement-contingent) and ’n’ to indicate ‘second stimulus’ (comparison stimulus). Making a response triggered a 750 ms inter-trial interval. If no response was made after 2500 ms, the next trial started automatically. Comparison stimuli were presented in a pseudorandom order where the same stimulus level was not shown consecutively more than twice.

Participants also completed active and passive motor control conditions, consisting of 60 trials each. The fixation cross, button press, visual and auditory stimuli, and inter-trial interval were identical to the experimental conditions. However, after the button press, there was a 1000 ms delay before the stimuli and question were presented. Active and passive trials were presented in mini blocks of 15 trials each, with the first 60 trials having the visual task and the second 60 having the auditory task.

### Stimuli

Stimuli were presented with Psychtoolbox (V 3.0.12) running on Octave (V 4.0.0) in Linux. Auditory stimuli consisted of a 1000 hz tone. The first tone was always presented at 74 dB, whereas the second had a loudness of 71, 72.5, 74, 75.5, or 77 dB. Visual stimuli consisted of a solid 250-pixel disc. The first disc was always presented at a luminance of 11.42 cd/m², whereas the second had a luminance of 8.84, 9.94, 11.42, 12.69, or 14.04 cd/m². Stimuli were presented for 50 ms. Luminance measurements were performed using an i1Display Pro photometer (X-Rite Pantone, Grand Rapids, USA). Volume measurements were performed using an RS-95 decibel metre (RS Components Ltd). The stimuli were presented on a fixed grey background with a luminance of 3.40 cd/m².

### Intensity Judgement Task

In the intensity judgement task, the proportion of ‘second brighter/louder’ responses was calculated and logistic psychometric functions were fitted using Psignifit 4 ^58^ for MATLAB, implementing a maximum-likelihood estimation, as described in Wichmann and Hill ^59^. Thresholds, representing the intensity value at which the stimulus of interest and the comparison stimulus were judged to be the same intensity in 50% of the trials (point of subjective equality; PSE) were derived for each participant and condition. As the luminance/loudness of the stimulus of interest was held constant throughout the experiment, a change in threshold across movement conditions would reflect purely perceptual differences. In this case, a shift towards lower intensities indicates that the comparison stimulus was judged as brighter or louder more often than the stimulus of interest ^for^ ^a^ ^comparable^ ^approach, see^ ^35^. We expected the perception of self-generated stimuli to be attenuated, and therefore we hypothesised that thresholds would be significantly lower in the active compared to the passive condition. Individual psychometric functions were visually inspected. For statistical analysis, the auditory and visual thresholds were entered separately into 2-tailed paired samples t-tests.

### Accuracy and Response Times

Accuracy was calculated by taking the proportion of correct responses per participant, per condition. Trials in which the stimulus of interest and comparison stimulus had identical intensity were not included in this analysis. Accuracies were entered into a 2×2 ANOVA with the factors of Modality (Visual, Auditory) and Movement (Active, Passive).

Response times were measured from the onset of the question and similarly entered into a 2×2 ANOVA.

### EEG Data Acquisition

EEG was continuously recorded at a sampling rate of 500 Hz from 32 active Ag/AgCl electrodes (Fp1/2, F7/8, F3/4, Fz, FT9/10, FC5/6, FC1/2, T7/8, C3/4, Cz, TP9/10, CP5/6, CP1/2, P7/8, P3/4, Pz, O1/2, and Oz). The EEG was referenced online to the electrode location FCz and the ground electrode was placed on the forehead. Impedances were kept at 25 kΩ or below. The signal was amplified by a BrainVision amplifier and recorded with BrainVision Recorder (Brain Products GmbH, Germany). Electrodes were mounted in an elastic cap (actiCAP, Brain Products GmbH, Germany) according to the international 10–20 system.

### EEG Preprocessing

Preprocessing was completed using the EEGLAB toolbox ^60^ in MATLAB (R2020a Mathworks, Sherborn, Massachusetts). EEG data were downsampled to 250 Hz and re-referenced to the average of electrodes TP9 and TP10. Line noise was removed using the Zapline Plus function ^61^. A high pass filter was applied at 1 Hz and the data were subjected to an extended infomax ICA. Components were classified using ICLabel ^62^ and any that were identified as muscle, eye or channel noise with greater than 79% estimated accuracy were removed. The ICA results were then applied to the unfiltered data. Finally, the data were bandpass filtered between 0.01 and 100 Hz.

Further analyses were completed with the Fieldtrip toolbox ^63^ and custom routines in MATLAB (R2020a Mathworks, Sherborn, Massachusetts). Trials in which the participant failed to make a response to the behavioural task and trials with unusually short button press response times (less than 100 ms) were excluded from all subsequent analyses. Further preprocessing steps specific to each analysis are described in the following sections (X, Y and Z).

### ERP- N1-P2 Peak-to-Peak

Data were segmented from 500 ms before to 1000 ms after the button press. Data were subjected to an artefact identification and rejection procedure, implemented with the ft_artifact_zvalue Fieldtrip function, which removes trials based on thresholding the z-transformed data. Segments containing data above or below 12 standard deviations from the mean of the z-transformed data, or above or below 9 standard deviations above or below the z-transformed data filtered between 100 and 120 Hz were removed. The latter criteria were used to detect muscle-based artefacts. The data were then bandpass filtered between 0.5 and 20 Hz. Next, the mean activity per participant, channel, and time point was calculated for each condition. The activity in the active and passive control conditions was then subtracted from the equivalent active/passive auditory and visual conditions. This method has been employed in previous experiments focusing on both auditory ^17,64^ and visual ^14,47,65,66^ ERPs.

Data were baseline corrected using the 100 ms interval between the button press and the stimulus presentation. Two regions of interest were defined for the ERPs. Due to the multimodal nature of the stimuli, we investigated electrodes typical of visual (O1, O2, Oz) and auditory (Cz, C3, C4) ERPs. ERPs were averaged separately across these two sets of electrodes for each of the four conditions (visual, active; visual, passive; auditory, active; auditory, passive). For statistical analysis, peak values were extracted by identifying the minimum (N1) or maximum (P2) value of the ERP across all conditions at the group level. A 24 ms time window was defined around this centre value. For visual electrodes, this procedure resulted in time windows of 80-104 ms for N1 and 132-156 ms for P2 and auditory electrodes, 76-100 ms for N1 and 212-236 ms for P2. The activity in these time windows was averaged together to produce a single value per condition, per participant. Comparable methods have been used in similar previous studies ^14,47,67^. In order to improve signal-to-noise ratio, we analysed and reported N1-P2 peak-to-peak amplitude, as in previous studies ^14,68^. N1-P2 peak-to-peak amplitude was calculated by subtracting the mean P2 peak amplitude from the mean N1 peak amplitude. The N1-P2 values were analysed for each of the two ROIs with 2×2 ANOVAs with the factors of movement (active, passive) and task modality (visual, auditory).

### Readiness Potential and Lateralized Readiness Potential

Data were segmented from 1300 ms before to 500 ms after the button press. Trial rejection was then performed with the ft_artifact_zvalue as described in section X (ERP- N1/P2). Data were then bandpass filtered between 0.01 and 40 Hz and baseline corrected between 1250 and 1000 ms before the button press. For RP analyses, data were averaged across electrodes Cz, C3, and C4. Lateralized readiness potential was calculated by subtracting the average of electrode C3 from the average of electrode C4. For statistical analysis, we did not have an a priori hypothesis regarding the exact time window(s) in which the conditions would differ. Therefore, we first tested the general difference between the visual and auditory conditions (averaged across active and passive) using a cluster-based permutation test. All time points were first tested with 2-tailed dependent samples t-tests with a significance level of p < .05. Contiguous time points exceeding this threshold were grouped into clusters and the sum of the t-values was used as the test statistic for the permutation test. This process was repeated 1000 times (1000 permutations) with shuffled condition labels, to determine a distribution of the probability of observing a cluster (or clusters) with that test statistic value. Clusters within the highest or lowest 2.5th percentile were considered significant.

The permutation tests revealed significant clusters beginning around 200 ms before the button press for RP. We based our second analysis on this time window. We calculated the mean amplitude across the significant clusters, resulting in one value per condition, per participant. We then subjected these values to a 2×2 repeated measures ANOVA with the factors of movement (active, passive) and task modality (visual, auditory).

### Multivariate pattern analysis with broad-band and band-specific EEG

We attempted to decode task modality with different sets of signals. Besides decoding based on the broad-band ERP data (preprocessed as in section EEG Preprocessing & ERP-N1/P2), we firstly applied a low-pass filter with 6Hz as high-cut off, aiming at retaining the low-frequency components that are distinguishable from the alpha and beta oscillations (see below, ^69^). Importantly, we did not apply an additional high-pass filter as in the univariate ERP analysis. Thus, the resulting signals potentially capture the ultra-slow RP and LRP components, which might subsume predictive information during movement preparation. We labeled these signals as the slow-ERP. We also focused on decoding based on narrow-band EEG within the alpha and beta bands. For this purpose, we Hilbert-transformed the band-filtered EEG and then extracted amplitudes and angles of these signals to obtain narrow-band amplitudes and instantaneous phase. We then conducted MVPA analysis based on these different signals (broad-band ERP, slow-ERP, alpha/beta amplitude and phase). We also carried out MVPA based on amplitude and phase within the delta and theta bands and reported the results in the supplement.

### Decoding across time, temporal generalisation, and cross-decoding

We then conducted a series of decoding analyses to decode task modality (visual vs. auditory). For all signals above, we first downsampled them to 50 Hz to reduce processing time. For decoding across time analyses (DAT), we trained a linear discriminant analysis (LDA) classifier ^70^ with a default hyperparameter lambda = 0.1 ^magnitude^ ^of^ ^shrinkage^ ^regularisation; 71^ at each time point, using all electrodes as features. This was done separately for the two movement conditions. To improve the signal-to-noise ratio, we sub-averaged all trials in groups of five trials, then cross-validated with 5-fold cross-validation (except for decoding based on instantaneous phase). This procedure was randomly repeated for 100 times. We used classifier accuracy at each time point as the performance metric. We then focused on the predefined time windows of interest within the motor preparation window (-0.4 to 0 seconds) and the stimulus perception window (0.2 to 0.5 seconds, corresponding to 0.1 to 0.4 seconds post-stimulus onset). We then compared the averaged decoding accuracy within both time windows, and compared them against chance (0.5) and against each other with one-sample and paired t-tests respectively. For illustrative purposes, we also statistically evaluate the significance of the decoding time series using a cluster-based permutation test (2-tailed dependent samples t-test, p-value threshold = .05, 10,000 permutations) at the group level, to identify clusters where the accuracy was significantly above chance (0.5). We also performed further cluster-based permutation tests to compare the decoding accuracies between the different task conditions. The tests had the same parameters, except that the test statistic underwent a Bonferroni correction for multiple comparisons (2 comparisons per condition, p = .05/2). We also conducted temporal generalisation analyses to evaluate the neural similarities between different time windows. To this end, the LDA classifier was trained to discriminate task modality at each time point and (t0) and tested on every other time point (t1), resulting in one 2-dimensional matrix of classification accuracies for each movement condition. All electrodes were included as features. The classification parameters were set identically to the DAT analysis. The statistical procedure was also analogous to that of the DAT analysis, except for the fact that the accuracy was averaged within a predefined 2D time window of interest, and the cluster-based permutation test was conducted also within a 2D matrix. At last, we conducted cross-decoding analyses to test the neural similarity between the active and passive conditions.

For this purpose, we trained LDA classifiers to decode task modality based on signals in the active condition for each time point. Then we tested the classifier performance on all other time points in the passive condition. This procedure was also implemented by training from the passive condition and testing on the active condition, and we averaged the decoding accuracy between these two steps. Except that no cross-validation is necessary for cross-decoding, all other classification parameters and statistical procedures were identical to temporal generalisation analysis.

## Acknowledgements

We thank Bianca van Kemenade and Daniel Reznik for valuable discussions on earlier versions of this manuscript. We also thank Katharina Schuster and Mona Rauschkolb for assistance with data collection and translation and Ruslan Spartakov and Julia Oppermann for assistance with data collection.

## Ethics

The study was approved by the local ethics committee of the medical faculty of the University of Marburg, Germany.

## AUTHOR CONTRIBUTIONS

Edward Ody: Conceptualization; data curation; formal analysis; investigation; methodology; software; validation; visualization; writing – original draft.

Tilo Kircher: Conceptualization; funding acquisition; project administration; resources; supervision; writing – review and editing.

Benjamin Straube: Conceptualization; funding acquisition; methodology; project administration; resources; supervision; writing – review and editing.

Yifei He: Conceptualization; formal analysis; methodology; supervision; writing – review and editing.

## References

1. Blakemore, S. J. & Frith, C. Self-awareness and action. Curr. Opin. Neurobiol. 13, 219– 224 (2003).

2. Wolpert, D. M. Computational approaches to motor control. Trends Cogn. Sci. 1, 209– 216 (1997).

3. Kilteni, K., Engeler, P. & Ehrsson, H. H. Efference Copy Is Necessary for the Attenuation of Self-Generated Touch. iScience 23, 100843 (2020).

4. Ford, J. M., Roach, B. J., Faustman, W. O. & Mathalon, D. H. Synch before you speak: Auditory hallucinations in schizophrenia. Am. J. Psychiatry 164, 458–466 (2007).

5. Cao, L., Thut, G. & Gross, J. The role of brain oscillations in predicting self-generated sounds. Neuroimage 147, 895–903 (2017).

6. Benedetto, A., Ho, H. T. & Morrone, M. C. The Readiness Potential Correlates with Action-Linked Modulation of Visual Accuracy. eNeuro 9, 1–9 (2022).

7. Vercillo, T., O’Neil, S. & Jiang, F. Action-effect contingency modulates the readiness potential. Neuroimage 183, 273–279 (2018).

8. Reznik, D., Simon, S. & Mukamel, R. Predicted sensory consequences of voluntary actions modulate amplitude of preceding readiness potentials. Neuropsychologia 119, 302–307 (2018).

9. Ody, E., Kircher, T., Straube, B. & He, Y. Pre-movement event-related potentials and multivariate pattern of EEG encode action outcome prediction. Hum. Brain Mapp. 44, 6198–6213 (2023).

10. Pinheiro, A. P., Schwartze, M., Gutiérrez-Domínguez, F. & Kotz, S. A. Real and imagined sensory feedback have comparable effects on action anticipation. Cortex 130, 290–301 (2020).

11. 11. Moran, C., Hogendoorn, H. & Landau, A. N. Neural evidence for action-related somatosensory predictions. *bioRxiv* (2024) doi:10.1101/2024.11.29.626056.

12. Martikainen, M. H., Kaneko, K.-I. & Hari, R. Suppressed responses to self-triggered sounds in the human auditory cortex. Cereb. Cortex 15, 299–302 (2005).

13. Ford, J. M., Palzes, V. A., Roach, B. J. & Mathalon, D. H. Did i do that? Abnormal predictive processes in schizophrenia when button pressing to deliver a tone. Schizophr. Bull. 40, 804–812 (2014).

14. Ody, E., Straube, B., He, Y. & Kircher, T. Perception of self-generated and externally-generated visual stimuli: Evidence from EEG and behavior. Psychophysiology (2023) doi:10.1111/psyp.14295.

15. Baess, P., Horváth, J., Jacobsen, T. & Schröger, E. Selective suppression of self-initiated sounds in an auditory stream: An ERP study. Psychophysiology 48, 1276–1283 (2011).

16. Blakemore, S. J., Wolpert, D. M. & Frith, C. D. Central cancellation of self-produced tickle sensation. Nat. Neurosci. 1, 635–640 (1998).

17. Bäß, P., Jacobsen, T. & Schröger, E. Suppression of the auditory N1 event-related potential component with unpredictable self-initiated tones: Evidence for internal forward models with dynamic stimulation. Int. J. Psychophysiol. 70, 137–143 (2008).

18. Sanmiguel, I., Todd, J. & Schröger, E. Sensory suppression effects to self-initiated sounds reflect the attenuation of the unspecific N1 component of the auditory ERP. Psychophysiology 50, 334–343 (2013).

19. Myers, J. C., Mock, J. R. & Golob, E. J. Sensorimotor Integration Can Enhance Auditory Perception. Sci. Rep. 16–18 (2020) doi:10.1038/s41598-020-58447-z.

20. Reznik, D., Henkin, Y., Schadel, N. & Mukamel, R. Lateralized enhancement of auditory cortex activity and increased sensitivity to self-generated sounds. Nat. Commun. 5, (2014).

21. van Kemenade, B. M., Arikan, B. E., Kircher, T. & Straube, B. Predicting the sensory consequences of one’s own action: First evidence for multisensory facilitation. Atten. Percept. Psychophys. 78, 2515–2526 (2016).

22. 22. Yon, D., Zainzinger, V., de Lange, F. P., Eimer, M. & Press, C. Action biases perceptual decisions toward expected outcomes. J. Exp. Psychol. Gen. 150, 1225–1236 (2021).

23. Haggard, P. & Whitford, B. Supplementary motor area provides an efferent signal for sensory suppression. Brain Res. Cogn. Brain Res. 19, 52–58 (2004).

24. Horváth, J. Action-related auditory ERP attenuation: Paradigms and hypotheses. Brain Res. 1626, 54–65 (2015).

25. Job, X. & Kilteni, K. Action does not enhance but attenuates predicted touch. Elife 12, e90912 (2023).

26. Kok, P., Jehee, J. F. M. & de Lange, F. P. Less is more: expectation sharpens representations in the primary visual cortex. Neuron 75, 265–270 (2012).

27. de Lange, F. P., Heilbron, M. & Kok, P. How Do Expectations Shape Perception?. Trends Cogn. Sci. 22, 764–779 (2018).

28. Press, C., Kok, P. & Yon, D. The Perceptual Prediction Paradox. Trends Cogn. Sci. 24, 13–24 (2020).

29. Yon, D., Gilbert, S. J., de Lange, F. P. & Press, C. Action sharpens sensory representations of expected outcomes. Nat. Commun. 9, 4288 (2018).

30. Saupe, K., Widmann, A., Bendixen, A., Müller, M. M. & Schröger, E. Effects of intermodal attention on the auditory steady-state response and the event-related potential. Psychophysiology 46, 321–327 (2009).

31. Alho, K., Woods, D. L., Algazi, A. & Näätänen, R. Intermodal selective attention. II. Effects of attentional load on processing of auditory and visual stimuli in central space. Electroencephalogr. Clin. Neurophysiol. 82, 356–368 (1992).

32. Karns, C. M. & Knight, R. T. Intermodal auditory, visual, and tactile attention modulates early stages of neural processing. J. Cogn. Neurosci. 21, 669–683 (2009).

33. Eimer, M. & Schröger, E. ERP effects of intermodal attention and cross-modal links in spatial attention. Psychophysiology 35, 313–327 (1998).

34. Blakemore, S. J., Wolpert, D. & Frith, C. Why can’t you tickle yourself? Neuroreport 11, R11–6 (2000).

35. Lubinus, C. et al. Action-based predictions affect visual perception, neural processing, and pupil size, regardless of temporal predictability. Neuroimage 263, 119601 (2022).

36. Ody, E., Kircher, T., He, Y. & Straube, B. Differential effects of self-initiated, externally triggered, and passive movements on action-outcome processing: Insights from sensory and motor-preparatory ERPs. (2023) doi:10.31219/osf.io/h279c.

37. Bae, G.-Y. & Luck, S. J. Decoding motion direction using the topography of sustained ERPs and alpha oscillations. Neuroimage 184, 242–255 (2019).

38 Hetenyi, D., Haarsma, J. & Kok, P. Pre-stimulus alpha oscillations encode stimulus-specific visual predictions. bioRxiv 2024.03. 13.584593 (2024) doi:10.1101/2024.03.13.584593.

39. Samaha, J., Boutonnet, B., Postle, B. R. & Lupyan, G. Effects of meaningfulness on perception: Alpha-band oscillations carry perceptual expectations and influence early visual responses. Sci. Rep. 8, 6606 (2018).

40. Limanowski, J., Litvak, V. & Friston, K. Cortical beta oscillations reflect the contextual gating of visual action feedback. Neuroimage 222, 117267 (2020).

41. King, J. R. & Dehaene, S. Characterizing the dynamics of mental representations: The temporal generalization method. Trends Cogn. Sci. 18, 203–210 (2014).

42. Brunia, C. H. M., van Boxtel, G. J. M. & Böcker, K. B. E. Negative Slow Waves as Indices of Anticipation: The Bereitschaftspotential, the Contingent Negative Variation, and the Stimulus-Preceding Negativity. The Oxford Handbook of Event-Related Potential Components 1–22 (2012) doi:10.1093/oxfordhb/9780195374148.013.0108.

43. Schmidt, S., Jo, H.-G., Wittmann, M. & Hinterberger, T. ‘Catching the waves’ - slow cortical potentials as moderator of voluntary action. Neurosci. Biobehav. Rev. 68, 639– 650 (2016).

44. Northoff, G. Slow cortical potentials and ‘inner time consciousness’ - A neuro- phenomenal hypothesis about the ‘width of present’. Int. J. Psychophysiol. 103, 174– 184 (2016).

45. Ford, J. M. & Mathalon, D. H. Electrophysiological evidence of corollary discharge dysfunction in schizophrenia during talking and thinking. J. Psychiatr. Res. 38, 37–46 (2004).

46. Hughes, G. & Waszak, F. ERP correlates of action effect prediction and visual sensory attenuation in voluntary action. Neuroimage 56, 1632–1640 (2011).

47. Mifsud, N. G. et al. Self-initiated actions result in suppressed auditory but amplified visual evoked components in healthy participants. Psychophysiology 53, 723–732 (2016).

48. Trajkovic, J., Di Gregorio, F., Thut, G. & Romei, V. Transcranial magnetic stimulation effects support an oscillatory model of ERP genesis. Curr. Biol. 34, 1048–1058.e4 (2024).

49. Arnal, L. H. & Giraud, A.-L. Cortical oscillations and sensory predictions. Trends Cogn.Sci. 16, 390–398 (2012).

50. Engel, A. K., Fries, P. & Singer, W. Dynamic predictions: oscillations and synchrony in top-down processing. Nat. Rev. Neurosci. 2, 704–716 (2001).

51. Mayer, A., Schwiedrzik, C. M., Wibral, M., Singer, W. & Melloni, L. Expecting to see a letter: Alpha oscillations as carriers of top-down sensory predictions. Cereb. Cortex 26, 3146–3160 (2016).

52. Roussel, C., Hughes, G. & Waszak, F. Action prediction modulates both neurophysiological and psychophysical indices of sensory attenuation. Front. Hum. Neurosci. 8, 115 (2014).

53. Schmalenbach, S. B., Billino, J., Kircher, T., van Kemenade, B. M. & Straube, B. Links between gestures and multisensory processing: Individual differences suggest a compensation mechanism. Front. Psychol. 8, 1–8 (2017).

54. Reznik, D., Henkin, Y., Levy, O. & Mukamel, R. Perceived loudness of self-generated sounds is differentially modified by expected sound intensity. PLoS One 10, 4–9 (2015).

55. Paraskevoudi, N. & SanMiguel, I. Self-generation and sound intensity interactively modulate perceptual bias, but not perceptual sensitivity. Sci. Rep. 11, 17103 (2021).

56. Reznik, D., Guttman, N., Buaron, B., Zion-Golumbic, E. & Mukamel, R. Action-locked Neural Responses in Auditory Cortex to Self-generated Sounds. Cereb. Cortex 31, 5560–5569 (2021).

57. World Medical Association. World Medical Association Declaration of Helsinki: ethical principles for medical research involving human subjects. JAMA 310, 2191–2194 (2013).

58. Schütt, H., Harmeling, S., Macke, J. & Wichmann, F. Psignifit 4: Pain-free Bayesian Inference for Psychometric Functions. J. Vis. 15, 474 (2015).

59. Wichmann, F. A. & Hill, N. J. The psychometric function: I. Fitting, sampling, and goodness of fit. Percept. Psychophys. 63, 1293–1313 (2001).

60. Delorme, A. & Makeig, S. EEGLAB: an open source toolbox for analysis of single-trial EEG dynamics including independent component analysis. J. Neurosci. Methods 134, 9–21 (2004).

61. Klug, M. & Kloosterman, N. A. Zapline-plus: A Zapline extension for automatic and adaptive removal of frequency-specific noise artifacts in M/EEG. Hum. Brain Mapp. 43, 2743–2758 (2022).

62. Pion-Tonachini, L., Kreutz-Delgado, K. & Makeig, S. ICLabel: An automated electroencephalographic independent component classifier, dataset, and website. Neuroimage 198, 181–197 (2019).

63. Oostenveld, R., Fries, P., Maris, E. & Schoffelen, J.-M. FieldTrip: Open source software for advanced analysis of MEG, EEG, and invasive electrophysiological data. Comput. Intell. Neurosci. 2011, 156869 (2011).

64. Whitford, T. J. et al. Electrophysiological and diffusion tensor imaging evidence of delayed corollary discharges in patients with schizophrenia. Psychol. Med. 41, 959–969 (2011).

65. 65. Csifcsák, G., et al. Action-associated modulation of visual event-related potentials evoked by abstract and ecological stimuli. Psychophysiology 56, e13289 (2019).

66. Mifsud, N. G. et al. Attenuation of visual evoked responses to hand and saccade-initiated flashes. Cognition 179, 14–22 (2018).

67. Saupe, K., Widmann, A., Trujillo-Barreto, N. J. & Schröger, E. Sensorial suppression of self-generated sounds and its dependence on attention. Int. J. Psychophysiol. 90, 300–310 (2013).

68. Timm, J., Schönwiesner, M., Schröger, E. & SanMiguel, I. Sensory suppression of brain responses to self-generated sounds is observed with and without the perception of agency. Cortex 80, 5–20 (2016).

69. Bae, G.-Y. & Luck, S. J. Dissociable Decoding of Spatial Attention and Working Memory from EEG Oscillations and Sustained Potentials. J. Neurosci. 38, 409–422 (2018).

70. McLachlan, G. J. Discriminant Analysis and Statistical Pattern Recognition. (John Wiley & Sons, 2005).

71. Ledoit, O. & Wolf M. Honey, I shrunk the sample covariance matrix. The Journal of Portfolio Management (2004).

