## Supplement for "Voluntary action sharpens sensory prediction and facilitates neural processing of contingent sensory stimuli"

### S1. Additional Behavioural Statistics

Table ST1. Behavioural Statistics

| | | $F(1,26)$ | $p$ | $\eta_p^2$ |
|---|---|---|---|---|
| Response Time | Movement | 1.7 | .206 | 0.06 |
| Response Time | Movement x Task | 0.01 | .928 | < 0.01 |
| Accuracy | Movement | 4.07 | .054 | 0.14 |
| Accuracy | Movement x Task | 0.38 | .543 | 0.02 |

### S2. Visual ERP post hoc tests

Table ST2. Visual ERP post-hoc test results.

| Comparison | | $t$ | $p_{bonf}$ |
|---|---|---|---|
| Visual, Active | Visual, Passive | -2.36 | .142 |
| Visual, Active | Auditory, Passive | 0.39 | >.999 |
| Auditory, Active | Auditory, Passive | 0.64 | >.999 |
| Auditory, Active | Visual, Passive | -2.03 | .289 |

### S3. Auditory Condition Confound

We previously observed a discrepancy between visual and auditory results in a contingent paradigm experiment using the same passive movement device [12]. As observed in the current data set, N1 amplitudes were larger in the active than the passive than the active condition, whereas P2 amplitudes showed the expected effect direction. Based on further analysis time-locked to the button press, which showed a negative peak resembling auditory N1 over the central electrodes for the passive movements regardless of task condition, we concluded that this unusual result was likely due to additional noise produced by the electromagnet. We speculated that the additional noise could have caused a sensory gating effect, masking the usual auditory attenuation in the active condition. Figure S1 shows a similar analysis in the current data, time-locked to the button press (time 0), averaged across electrodes Cz, C3 and C4 and baseline-corrected between -100 to 0 ms before the button press.

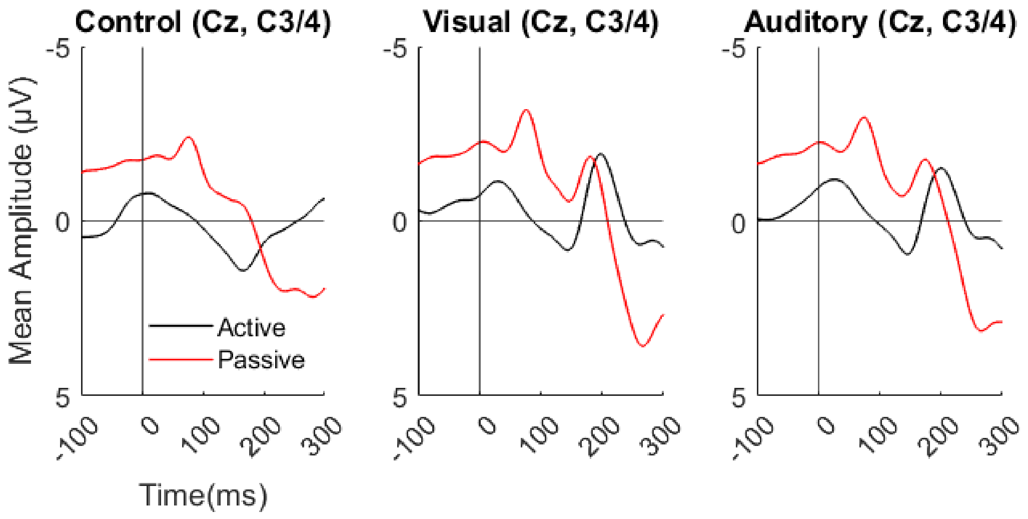

Figure S1. Average ERPs at C3/4 and Cz, time locked to button press (0) for control, visual and auditory conditions.

### S4. Additional ERP results

In the Auditory sensory ERPs, the effect of movement was significant ($F(1, 27)$ = 11.25, $p$ = .002, $\eta_p^2$ = 0.3), with higher N1-P2 peak-to-peak amplitudes in the active (EMM = 5.49 μV) than passive (EMM = 3.75 μV). The interaction between Task and Movement was also significant ($F(1, 27)$ = 7.27, $p$ = .012, $\eta_p^2$ = 0.21). Bonferroni-corrected post hoc tests showed a significant difference between active and passive for the visual task condition (t = 3.16, $p$ = .023). The difference between active and passive for the auditory condition was not significant (t = -0.1, $p$ > .999). Additional comparisons are presented in Table STX. The effect of Task was not significant ($F(1,27)$ = 1.95, $p$ = .174, $\eta_p^2$ = 0.07).

Table ST3. Auditory ERP post-hoc test results.

| Comparison | | *t* | $p_{bonf}$ |
|---|---|---|---|
| Visual, Active | Visual, Passive | 4 | .003** |
| Visual, Active | Auditory, Passive | 3.32 | .015** |

| | | | |
|---|---|---|---|
| Auditory, Active | Auditory, Passive | 1.72 | .586 |
| Auditory, Active | Visual, Passive | 1.42 | >.999 |

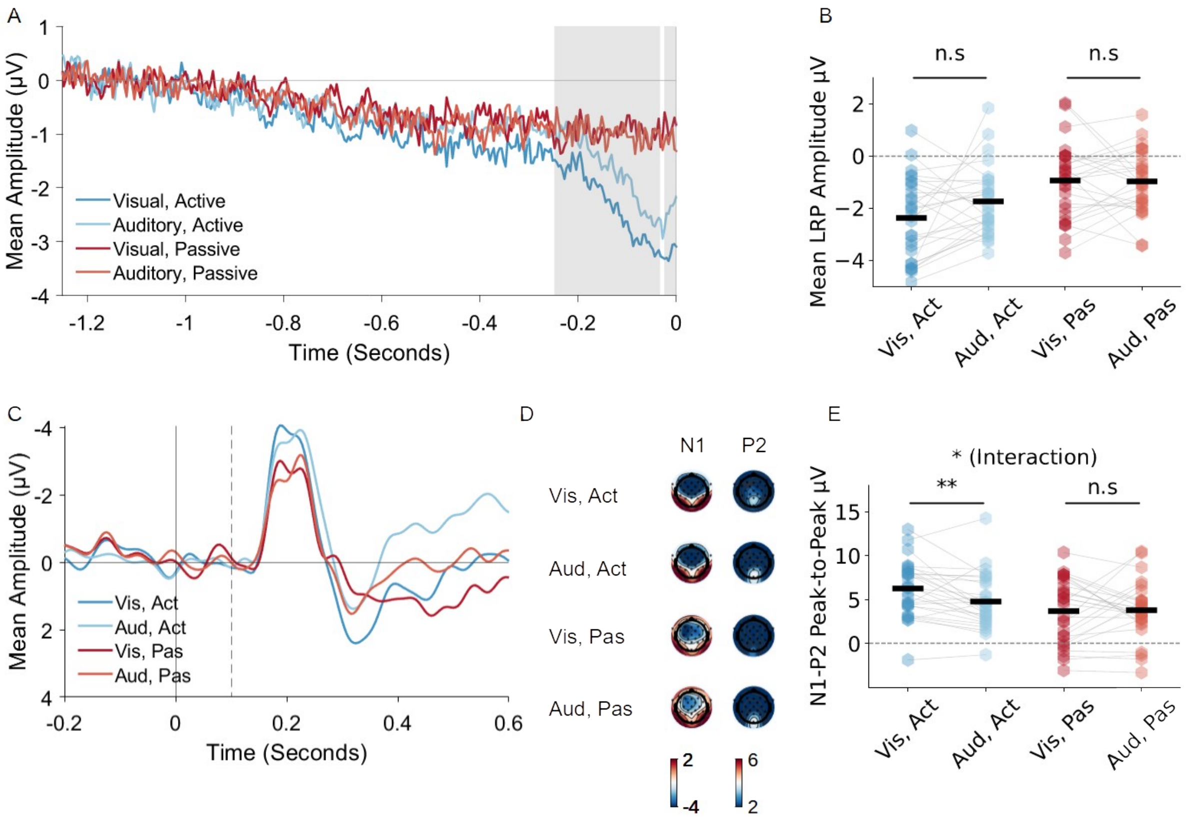

Figure S2. Event-related potential results. (A): Lateralized readiness potential. Average of electrode C3 - electrode C4, time-locked to the button press (t = 0). (B) Mean amplitudes averaged across all significant time points identified in the permutation test. (C) Topographical plots for all experimental conditions in 200 ms bins for the 1200 ms time period preceding the button press. (D) Visual ERPs averaged across electrodes Oz, O1, and O2, time-locked to the button press (0), with stimulus presentation at 0.1 seconds (dashed line). Baseline period 0 to 0.1 seconds (interval between button press and stimulus). (E) Topographical plots for the N1 (X-Y ms) and P2 (X-Y ms) peaks. (F) Mean N1-P2 peak-to-peak amplitude for the four conditions, $p < .001$: ***, $p < .01$: **. $p < .05$: * (Bonferroni-corrected).

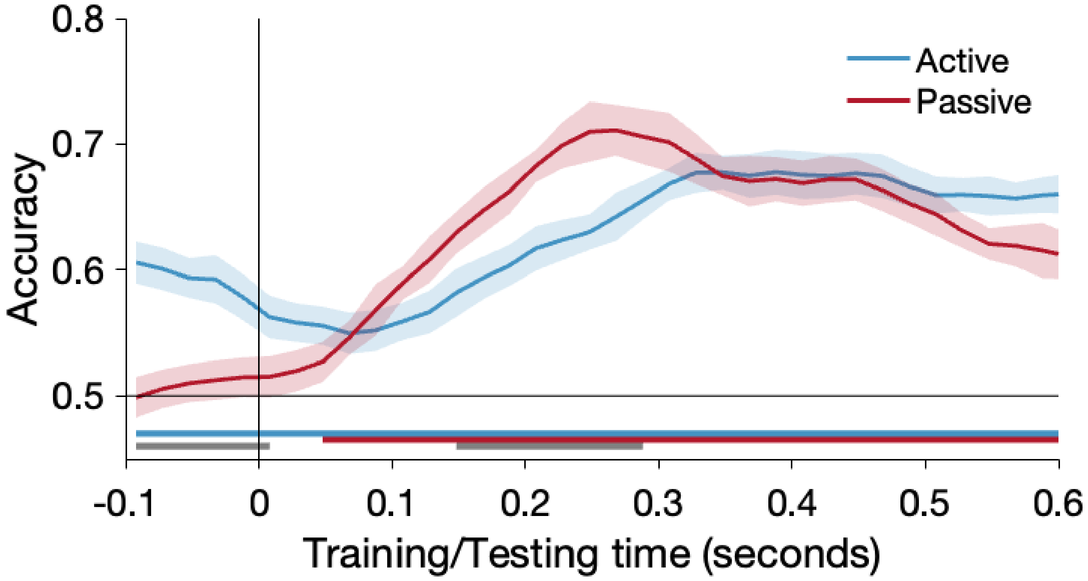

Figure S3. Decoding accuracy time series (means and standard errors) for the active (blue) and the passive (red) conditions based on the broadband ERPs, baseline correction based on the 0 to 0.1 seconds time window after the button press (-0.1 to 0 seconds before the onset of the audiovisual stimulus). Here, despite application of a different baseline, the decoding difference between the passive and active condition remains significant (higher decoding for the passive condition). Horizontal thick lines of respective colours indicate the time windows where the accuracy of both conditions was significantly above chance (cluster $p < .05$, corrected). The grey horizontal thick line illustrates the time windows where the accuracy of both conditions significantly differ (cluster $p < .05$, corrected). Time zero marks the onset of the button press. Audiovisual stimulus onsets at 0.1 seconds.

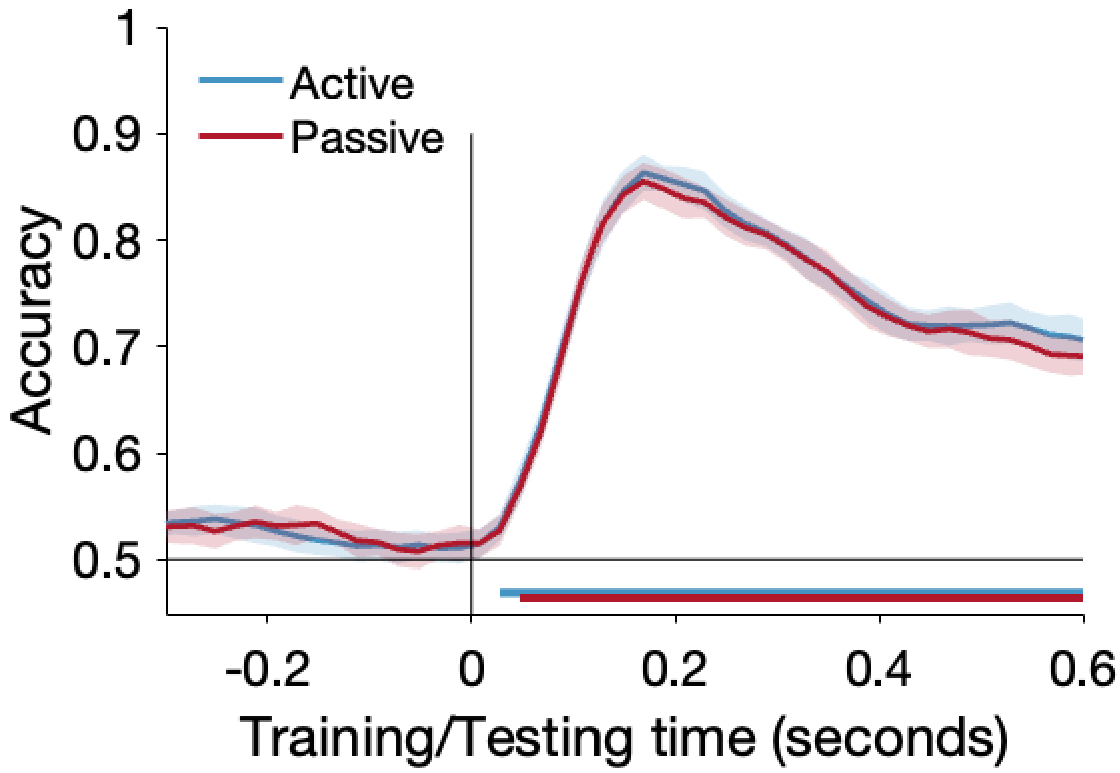

Figure S4. Decoding accuracy time series (means and standard errors) for the active (blue) and the passive (red) conditions based on the broadband ERPs, time-locked to the onset of the second visual or auditory stimulus (baseline -0.1 to 0 seconds). Here, despite the generally high accuracy for decoding two unimodal sensory signals, we observed no difference between the decoding time series for active vs. passive condition. Horizontal thick lines of respective colours indicate the time windows where the accuracy of both conditions was significantly above chance (cluster $p < .05$, corrected). The grey horizontal thick line illustrates the time windows where the accuracy of both conditions significantly differ (cluster $p < .05$, corrected). Time zero marks the onset of the button press. Audiovisual stimulus onsets at 0.1 seconds.

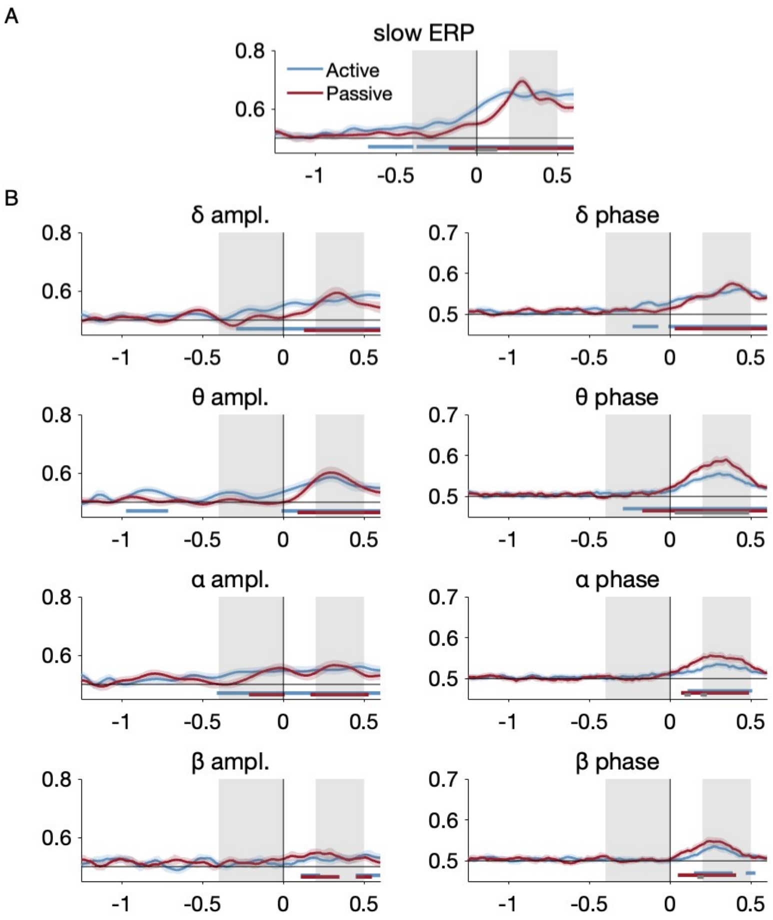

Figure S5. Decoding across time accuracy time series (means and standard errors) based on the (A) slow ERP and (B) amplitude and instantaneous phase within the delta, theta, alpha, and beta bands to decode task modality for the active (blue) and the passive (red) conditions. For all panels, horizontal thick lines of respective colours indicate the time windows where the accuracy of both conditions was significantly above chance (cluster $p < .05$, corrected). The grey horizontal thick line illustrates the time windows where the accuracy of both conditions significantly differ (cluster $p < .05$, corrected). Time zero marks the onset of the button press. Audiovisual stimulus onsets at 0.1 seconds. Light grey patches highlight the time windows of interest (TOI) for motor preparation (prep., -0.4 to 0 seconds) and for stimulus perception (stim., 0.2 to 0.5 seconds, corresponds to 0.1 to 0.4 seconds after stimulus onset).

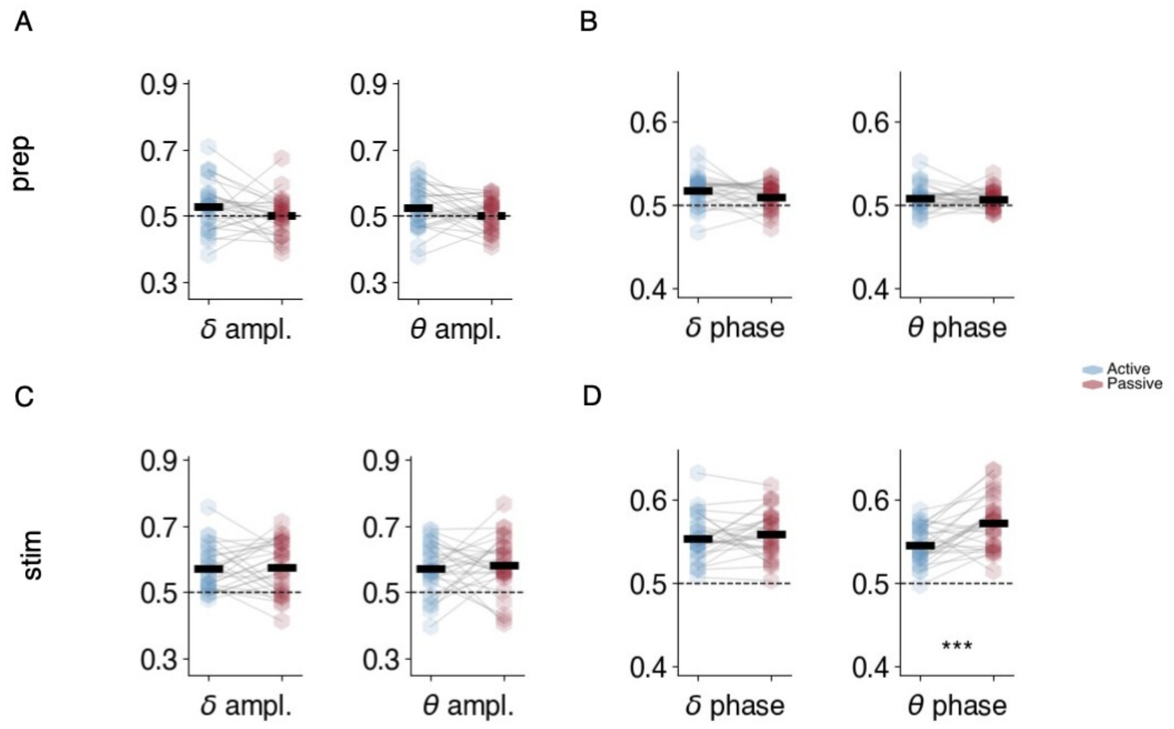

Figure S6. (A&B): Averaged decoding accuracy based on the delta and theta amplitude and phase within the motor preparation window (prep., -0.4 to 0 seconds). Here we observed no significant decoding difference between the movements. (C&D): Averaged decoding accuracy based on the delta and theta amplitude and phase within the stimulus window  (stim., 0.2 to 0.5 seconds, corresponds to 0.1 to 0.4 seconds after stimulus onset). Significant decoding difference between movements was only observed in the theta phase (D, left), suggesting its phase-locking nature. For all panels, horizontal black bars indicate the mean across all participants. The means significantly above chance are marked as solid; when they do not differ significantly from chance they are marked as dotted. Asterisks indicate statistical significance of dependent-sample t-tests between the active and passive conditions, $p < .001$: ***, $p < .01$: **. $p < .05$: *.

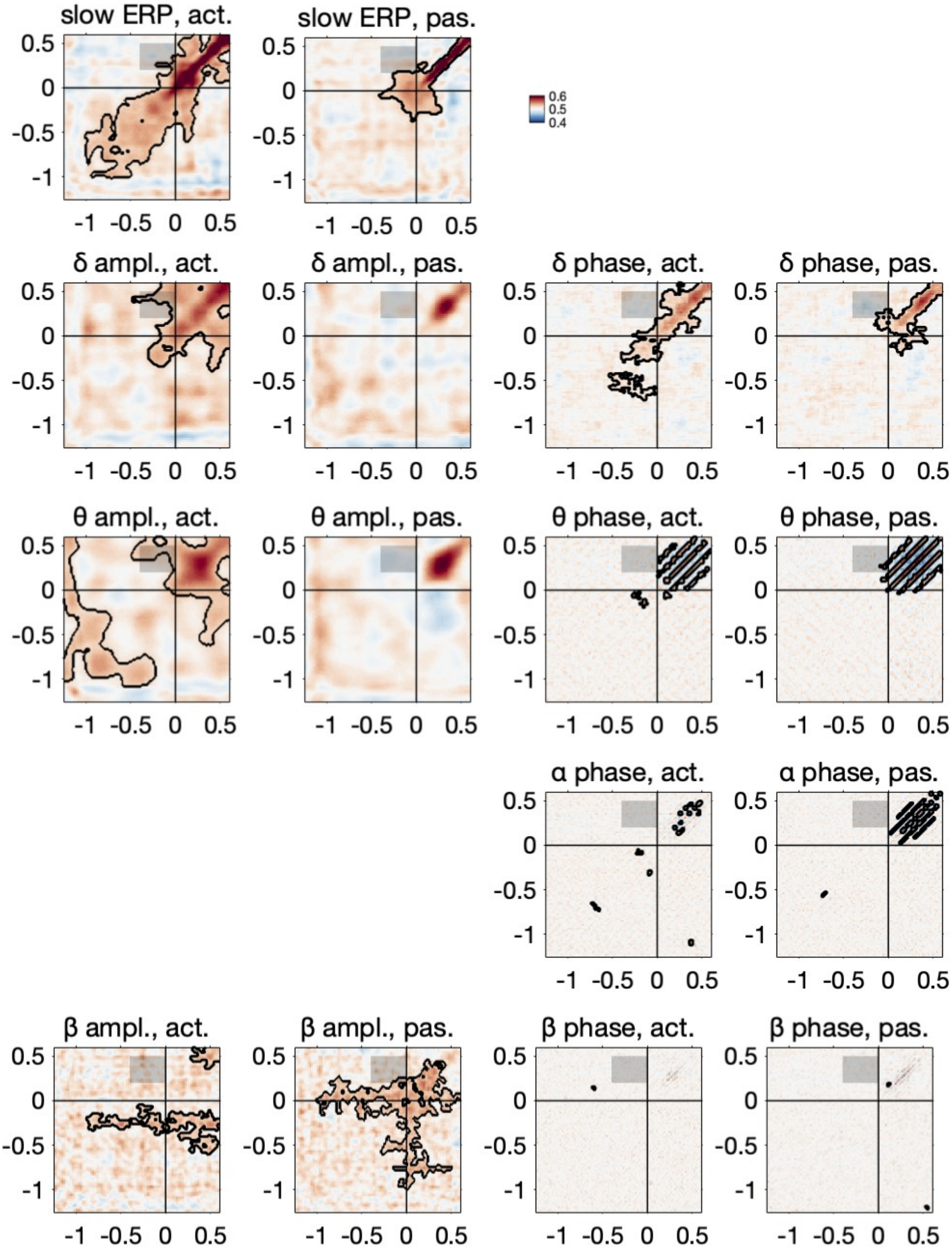

Figure S7. Temporal generalisation based on the slow ERPs, the amplitudes within the delta, theta, and beta bands, and phase signals from delta, theta, alpha, and beta bands, for the active and for passive conditions. Time windows of interest (training from the motor preparation window (-0.4 to 0 seconds) and testing on the stimulus perception window (0.2 to 0.5 seconds, corresponds to 0.1 to 0.4 seconds after stimulus onset) is highlighted in the light gray patch. Black contour indicates training-testing time point combinations with above-chance decoding, corrected with the cluster-based permutation tests.

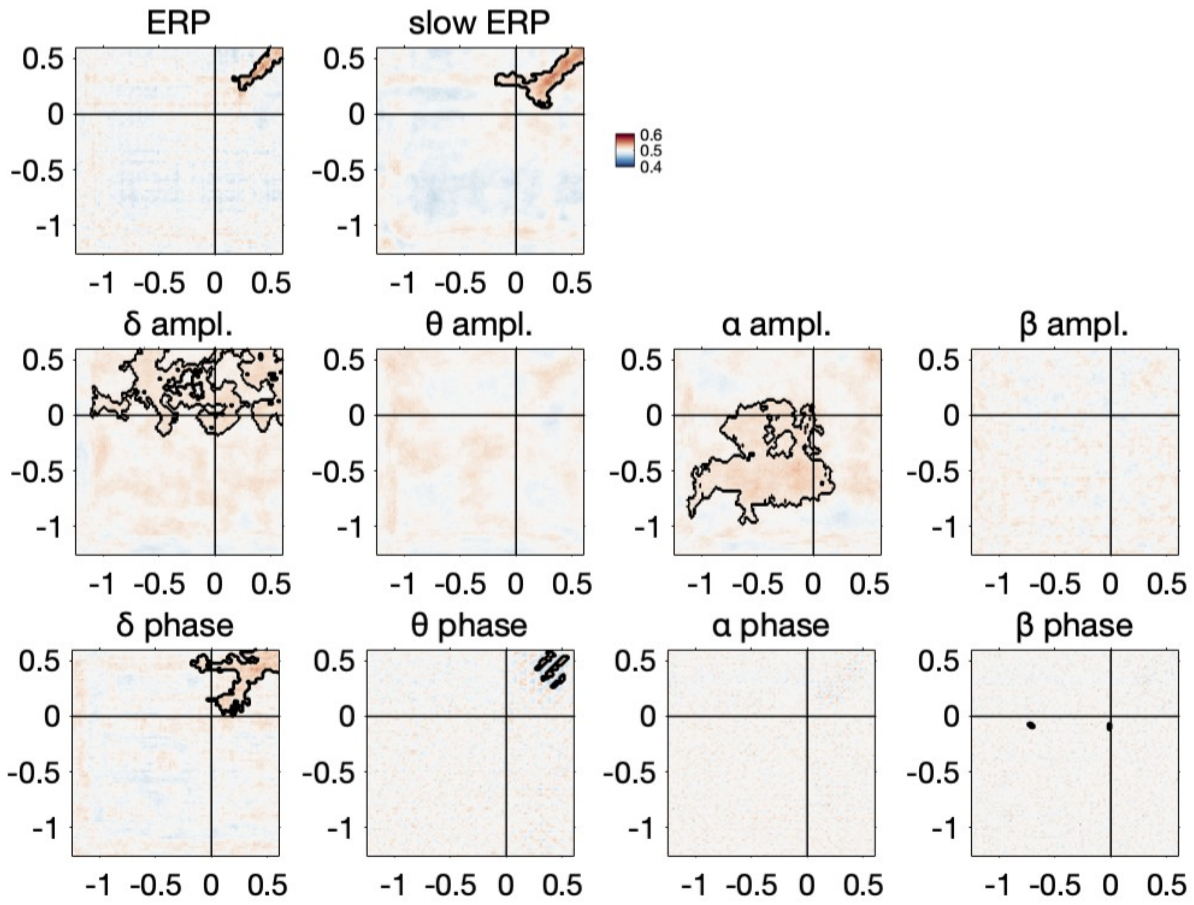

Figure S8. Cross-decoding results between the active and the passive conditions. The broadband ERP and the slow ERP suggest neural similarity between movements within the stimulus perception but not the motor preparation (top). Regarding the amplitudes within the delta, theta, alpha, and beta bands (middle), most importantly, the alpha amplitude carries comparable pattern between movements only during motor preparation. For the instantaneous phase within the delta, theta, alpha and beta bands (bottom), neural similarity between movements was also only observed during stimulus perception. For all sub-figures, black contour indicates training-testing time point combinations with above-chance decoding, corrected with the cluster-based permutation tests.
